# A pan-variant miniprotein inhibitor protects against SARS-CoV-2 variants

**DOI:** 10.1101/2024.08.08.606885

**Authors:** Jimin Lee, James Brett Case, Young-Jun Park, Rashmi Ravichandran, Daniel Asarnow, M. Alejandra Tortorici, Jack T. Brown, Shilpa Sanapala, Lauren Carter, David Baker, Michael S. Diamond, David Veesler

**Author notes:** These authors contributed equally (Jimin Lee, James Brett Case).

## Abstract

The continued evolution of severe acute respiratory syndrome coronavirus 2 (SARS-CoV-2) has compromised neutralizing antibody responses elicited by prior infection or vaccination and abolished the utility of most monoclonal antibody therapeutics. We previously described a computationally-designed, homotrimeric miniprotein inhibitor, designated TRI2-2, that protects mice against pre-Omicron SARS-CoV-2 variants. Here, we show that TRI2-2 exhibits pan neutralization of variants that evolved during the 4.5 years since the emergence of SARS-CoV-2 and protects mice against BQ.1.1, XBB.1.5 and BA.2.86 challenge when administered post-exposure by an intranasal route. The resistance of TRI2-2 to viral escape and its direct delivery to the upper airways rationalize a path toward clinical advancement.

## Introduction

The severe acute respiratory syndrome coronavirus 2 (SARS-CoV-2) spike (S) glycoprotein interacts with its host receptor ACE2 and initiates viral entry into cells^1–4^. The emergence of SARS-CoV-2 Omicron variants at the end of 2021 and afterwards have reduced the efficacy of vaccines and monoclonal antibodies, increased the number of reinfections or breakthrough infections, and led to successive waves of global infection^5–10^.

We previously described a computationally-designed, homotrimeric miniprotein inhibitor, designated TRI2-2, that binds with high avidity to SARS-CoV-2 S as a result of simultaneously engaging all three receptor-binding domains (RBDs) within a S trimer^11,12^. We showed that intranasal administration of TRI2-2 after viral exposure protected mice from challenge with the SARS-CoV-2 Beta and Delta variants^12^. The low manufacturing cost of TRI2-2 along with its antiviral efficacy when delivered to the upper respiratory tract are attractive properties for the development of next-generation countermeasures blocking SARS-CoV-2 at the site of initial infection.

## Results

To investigate the ability of TRI2-2 to recognize SARS-CoV-2 Omicron variants associated with recent infection waves, we assessed binding to a panel of biotinylated RBDs immobilized on biolayer interferometry biosensors. TRI2-2 bound with single digit picomolar avidities to the Wuhan-Hu-1 and Delta RBDs and with nanomolar avidities to the BA.1, BA.2, BA.2.12.1, BA.2.75.2, BA.5, BQ.1.1, XBB.1.5, BA.2.86, and JN.1 RBDs (**Fig. 1A and Fig S1**). These data establish that TRI2-2 binds avidly (i.e. with slow off-rates) to all Omicron variants evaluated despite accumulation of RBD mutations in the receptor-binding motif (ACE2-binding site).

**Figure 1.**
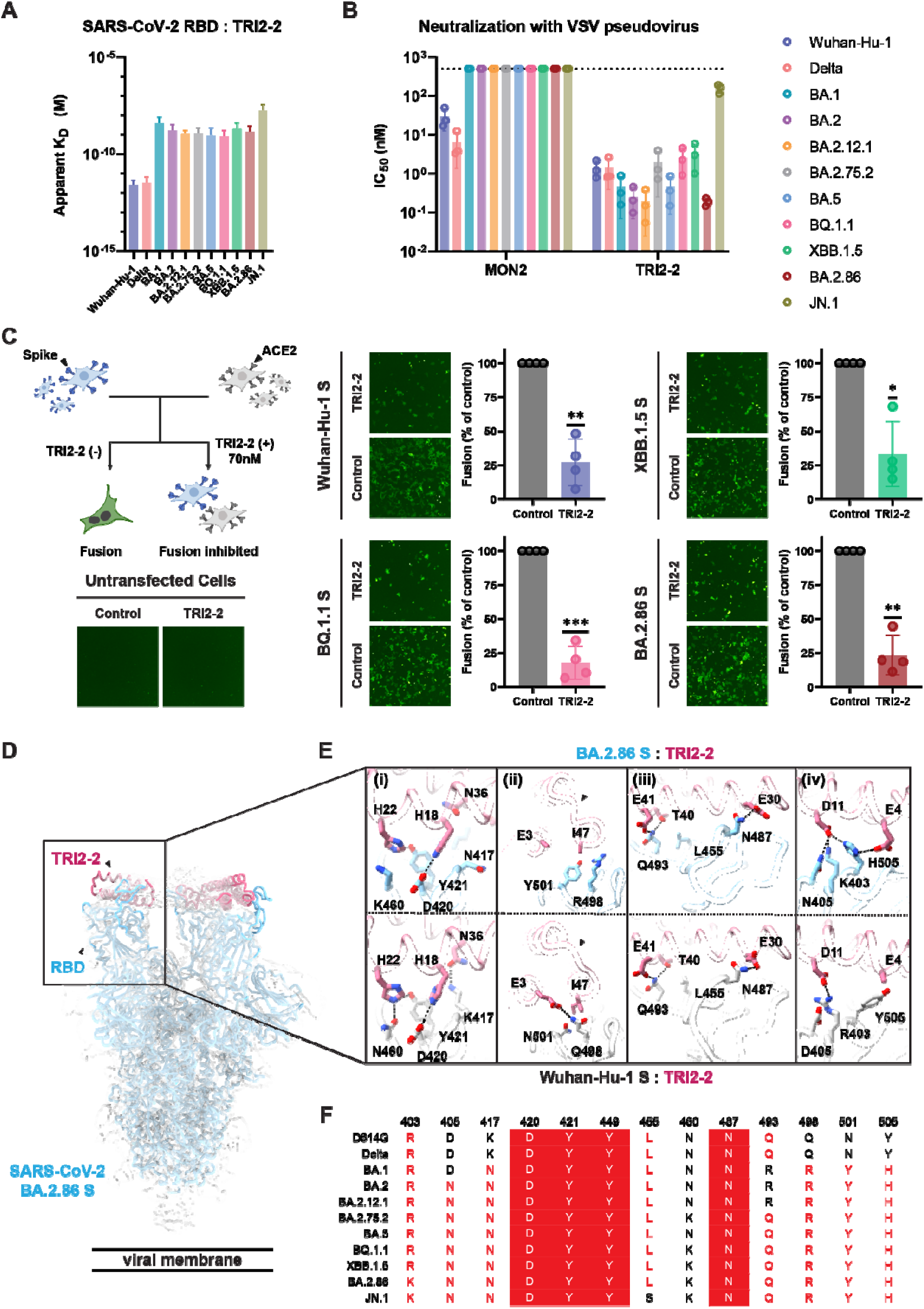
TRI2-2 cross-reacts with and potently neutralizes SARS-CoV-2 Omicron variants. **(A)** Binding of TRI2-2 to variant RBDs immobilized at the surface of BLI biosensors. Means of two biological replicates are shown as bar graphs with lines representing SD. **(B)** Neutralization of SARS-CoV-2 S VSV pseudoviruses harboring the Wuhan-Hu-1 D614G, Delta, BA.1, BA.2, BA.2.12.1, BA.2.75.2, BA.5, BQ.1.1, XBB.1.5, BA.2.86, or JN.1 S. Means of three biological replicates (each replicate is shown as a circle) rendered as bar graphs with SD. **(C)** Cell-cell fusion assay between BHK21 cells expressing SARS-CoV-2 D614G, BQ.1.1, XBB.1.5, or BA.2.86 S glycoprotein and VeroE6-TMPRSS2 cells in the absence of TRI2-2 (control) or in the presence of 70nM TRI2-2. Each dot represents replicate from four different biological replicates. SDs shown as lines. One sample t tests between control and TRI2-2 treatment; ns, not significant; *P□<□0.05, **P□<□0.01, ***P□<□0.001, ****P□<□0.0001). The schematic of the cell-cell fusion assay was rendered using Biorender. **(D-E)** CryoEM structure of TRI2-2 bound to the BA.2.86 S glycoprotein trimer (electron potential map shown as a semi-transparent gray surface) (D) and close-up views of the binding interface between TRI2-2 and BA.2.86 RBD compared to that obtained in complex with the Wuhan-Hu-1 RBD (PDB 7UHB) (E). **(F)** Amino acid sequences of the key residues at the binding interface. Coloring scheme follows ESPript 3^18^.

We subsequently tested the capacity of TRI2-2 for inhibition of vesicular stomatitis virus (VSV) particles pseudotyped with the Wuhan-Hu-1 D614G, Delta, BA.1, BA.2, BA.2.12.1, BA.2.75.2, BA.5, BQ.1.1, XBB.1.5, BA.2.86 or the JN.1 S using HEK293T target cells stably expressing human ACE2^13^. TRI2-2 potently neutralized all pseudoviruses tested, in a concentration-dependent manner, with half-maximal inhibition concentrations ranging between ∼0.5 and ∼5 nM across all variants except for JN.1, for which its potency was reduced to ∼300 nM (**Fig. 1B and Fig S2**). Comparatively, the monomeric AHB2 minibinder, which is the module TRI2-2 is built from (through genetic fusion to a trimerization motif), inhibited Wuhan-Hu-1 D614G S- and Delta S VSV pseudoviruses but failed to block any of the Omicron variants evaluated (**Fig. 1B and Fig S2**). These findings demonstrate that harnessing the binding avidity resulting from trivalent engagement of S trimers endows TRI2-2 with variant-resistant neutralizing activity.

We subsequently assessed the ability of TRI2-2 to inhibit syncytia formation between cells using a split green fluorescent protein (GFP) system with VeroE6/TMPRSS2 target cells (VeroE6 cells stably expressing TMPRSS2 and GFP β strand 11) and BHK-21 effector cells (stably expressing GFP β strands 1 to 10) transiently transfected with Wuhan-Hu-1 D614G, BQ.1.1, XBB.1.5 or BA.2.86 S^6,14^. Consistent with the pan-variant neutralizing activity observed, addition of TRI2-2 dampened S-mediated syncytia formation with all variants tested (**Fig. 1C and Fig S3**)^15^.

To determine the structural basis for the observed resilience to antigenic changes in the S glycoprotein of SARS-CoV-2 variants, we determined a cryoEM structure of the BA.2.86 S trimer bound to TRI2-2 at 2.4 Å resolution applying C3 symmetry (**Fig. 1D and Fig S4**). Local refinement of the region comprising the RBD and TRI2-2 yielded a reconstruction at 3.2 Å resolution with improved resolvability, revealing key interacting residues that are mutated or conserved in the SARS-CoV-2 Wuhan-Hu-1 RBD and BA.2.86 RBD (**Fig. 1E**). The BA.2.86 S K417N_SARS-CoV-2_ **(Fig.1E-i)** and Q498R_SARS-CoV-2_ **(Fig.1E-ii)** mutations abolish hydrogen bonds formed with N36_TRI2-2_ and E3_TRI2-2_, respectively, possibly participating in reducing the binding avidity observed for Omicron variants relative to SARS-CoV-2 Wuhan-Hu-1 and Delta. The Q498R and N501Y mutations are sterically incompatible with the positioning of the TRI2-2 I47 side chain observed in the Wuhan-Hu-1 complex structure and leads to reorganization of the minibinder region comprising the C-terminal part of the second helix and loop connecting to the third helix **(Fig.1E-ii)**. Furthermore, our structure shows that the JN.1 L455S_SARS-CoV-2_ **(Fig.1E-iii)** residue substitution reduces van der Waals packing at the interface with the minibinder, thereby dampening TRI2-2 binding avidity and consequently neutralizing activity. However, hydrogen bonds formed between H18_TRI2-2_ and D420_SARS-CoV-2_ **(Fig.1E-i)**, H22_TRI2-2_ and Y421_SARS-CoV-2_ **(Fig.1E-i)**, E30_TRI2-2_ and N487_SARS-CoV-2_ **(Fig.1E-iii)**, T40_TRI2-2_/E41_TRI2-2_ and Q493_SARS-CoV-2_ **(Fig.1E-iii)**, are conserved in the Wuhan-Hu-1 and BA.2.86 RBD complex structures. We further note that although BA.1 and BA.2 harbored the Q493R substitution^6,7^, TRI2-2 retained potent neutralizing activity against these variants (**Fig. 1F**). Contacts in several regions are remodeled by residue changes between Wuhan-Hu-1 and BA.2.86, as revealed by our structural data. The BA.2.86 S Y505H substitution leads to formation of a salt bridge triad with TRI2-2 residues E4 and D11, while D11 is salt bridged and hydrogen bonded to K403 and N405 (which are mutated from R403 and D405 in Wuhan-Hu-1 S), respectively **(Fig.1E-iv)**. Moreover, the BA.2.86 S N460K_SARS-CoV-2_ mutation replaces the hydrogen bond formed with H22_TRI2-2_ by a cation-pi interaction with the same residue **(Fig.1E-i)**. Overall, TRI2-2 buries a comparable surface at the interface with the Wuhan-Hu-1 and the BA.2.86 RBDs despite the aforementioned residue mutations. These data unveil the molecular basis for retained TRI2-2 binding and neutralization of SARS-CoV-2 variants that have emerged over the past 4.5 years despite variations within the targeted epitope, underscoring its exceptional resilience to antigenic changes.

To study the protective efficacy of TRI2-2 *in vivo* against immune evasive SARS-CoV-2 Omicron variants, we intranasally inoculated highly susceptible K18-hACE2 mice^16^ with 10^4^ FFU of BQ.1.1, XBB.1.5, or BA.2.86. One day later, we intranasally administered a single 10 mg/kg dose of TRI2-2 or an influenza virus control minibinder^17^. For all variants evaluated, post-exposure TRI2-2 treatment protected against weight loss throughout the duration of the experiments and reduced viral titers in the lungs and nasal turbinates six days post-challenge as compared to the control (influenza virus) minibinder (**Fig. 2 and Fig S5**). These results indicate that intranasal administration of TRI2-2 confers protection against SARS-CoV-2 challenge in a stringent model of disease with three key SARS-CoV-2 Omicron variants.

**Figure 2.**
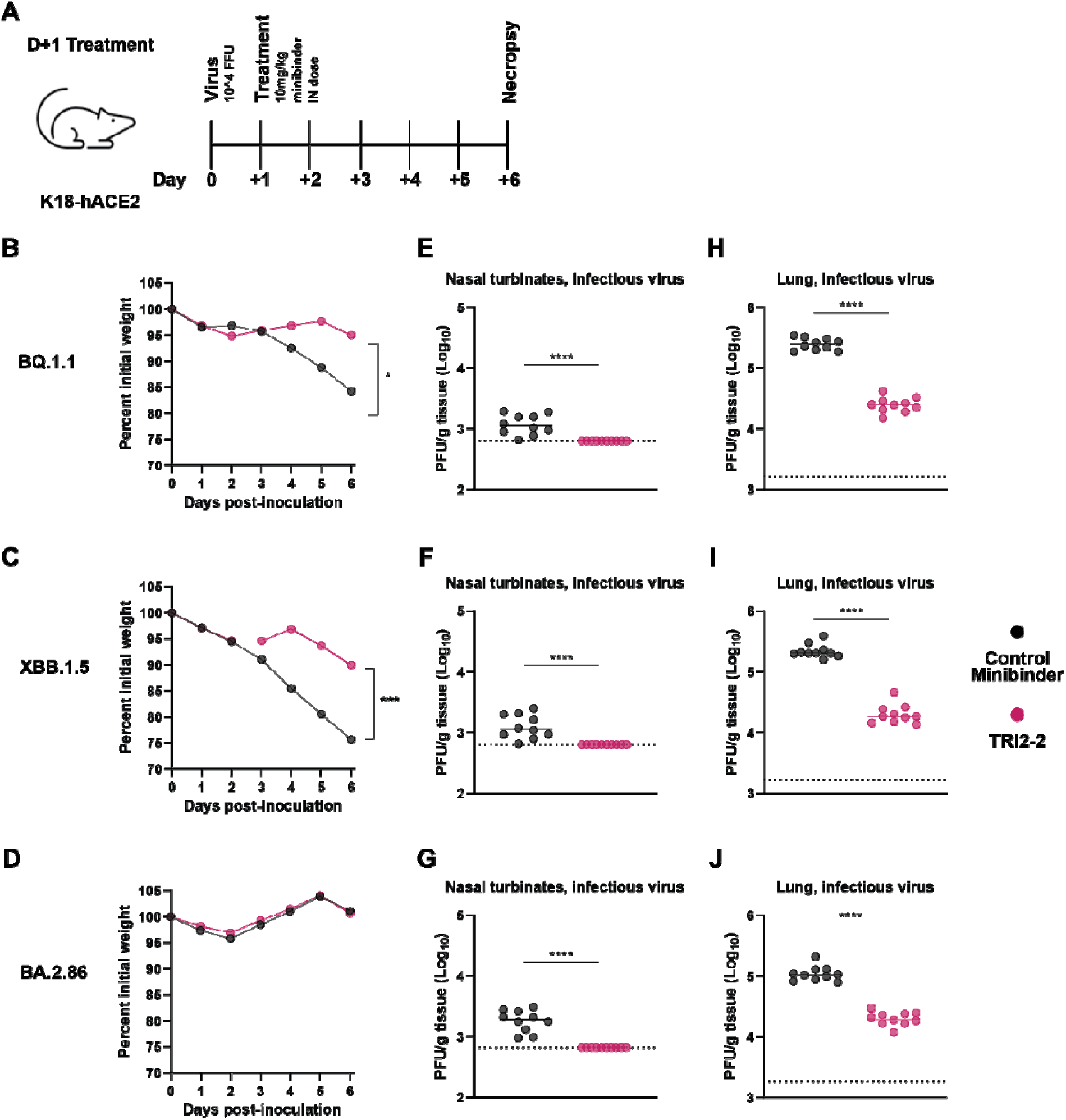
Post-exposure TRI2-2 administration protects mice challenged with the SARS-CoV-2 BQ.1.1, XBB.1.5, and BA.2.86 variants. **(A)** Schematic of study design. **(B-D)** Weight loss for mice challenged with BQ.1. (B), XBB.1.5 (C), or BA.2.86 (D) (dots represent mean.; n=10 mice per group per challenge virus; differences in area under the curves assessed by Student’s t-test with Welch’s correction for each virus; *P < 0.01; ***P<0.001). **(E-G)** Nasal turbinate infectious viral titers for mice challenged with BQ.1.1 (E), XBB.1.5 (F), or BA.2.86 (G). **(H-J)** Lung infectious viral titers for mice challenged with BQ.1.1 (H), XBB.1.5 (I), or BA.2.86 (J) (lines indicate median.; n=10 mice per group per challenge virus, two experiments; Two-tailed Mann-Whitney test between control and TRI2-treatment; ns, not significant; *P□<□0.05, ***P□<□0.001, ****P□<□0.0001.).

## Discussion

SARS-CoV-2 breakthrough infections elicit more robust neutralizing antibody titers in the human upper respiratory tract at the site of initial infection, than intramuscular vaccination alone^19–21^. These findings, along with waning of binding and neutralizing antibodies, likely contribute to the continued transmission of SARS-CoV-2 globally and motivate the development and evaluation of next-generation countermeasures that may be administered intranasally or orally. Preclinical assessment of intranasally administered influenza and sarbecovirus vaccine candidates demonstrated the induction of lung-resident protective mucosal humoral and cellular immunity at the site of viral entry^22–26^ and lipopeptide fusion inhibitors prevented SARS-CoV-2 direct-contact transmission in ferrets^27^. Furthermore, post-exposure prophylaxis nasal spray administration of the SA58 monoclonal antibody in humans was shown to markedly reduce the risk of contracting COVID-19^28^.

The computationally-designed TRI2-2 minibinder mediates pan-variant neutralizing activity and *in viv*o protection of mice in both the upper and lower airways against the highly immune evasive SARS-CoV-2 BQ.1.1, XBB.1.5, and BA.2.86 variants. These data show that TRI2-2 can accommodate residue substitutions within its epitope and provide a molecular framework to explain the remarkable retained neutralization of variants that have emerged since the pandemic began in 2019. Moreover, TRI2-2 is endowed with exceptional biophysical stability, enabling cost-effective, large-scale microbial production, setting it apart from monoclonal antibodies that are expensive to manufacture and more challenging to scale. TRI2-2 will be evaluated in humans in an upcoming clinical trial and could herald a new era of computationally-designed prophylactics and therapeutics.

## Author Contributions

J.L., J.B.C., M.S.D., and D.V. designed the experiments; R.R. recombinantly expressed and purified TRI2-2. J.L. performed binding assays and neutralization assays. J.L. carried out fusion assays with help from M.A.T.. J.L. vitrified the specimen and carried out cryoEM data collection. J.L, and Y.J.P., processed the cryoEM data with help from D.A. and D.V.. J.L and D.V. built and refined the atomic models. J.B.C. carried out the mice challenge study with assistance from S.S.. J.L., J.B.C., and D.V. analyzed the data and wrote the manuscript with input from all authors; D.B., M.S.D., and D.V. supervised the project.

## Acknowledgements

This study was supported by the National Institute of Allergy and Infectious Diseases (R01AI160052 to D.B. and D.V., R01 AI157155 and P01 AI168347 to M.S.D., DP1AI158186 and 75N93022C00036 to D.V.), a Pew Biomedical Scholars Award (D.V.), an Investigators in the Pathogenesis of Infectious Disease Awards from the Burroughs Wellcome Fund (D.V.), the University of Washington Arnold and Mabel Beckman cryoEM center and the National Institute of Health grant S10OD032290 (to D.V.). D.B. and D.V. are Investigators of the Howard Hughes Medical Institute and D.V. is the Hans Neurath Endowed Chair in Biochemistry at the University of Washington.

---

**Supplementary Figure 1.**
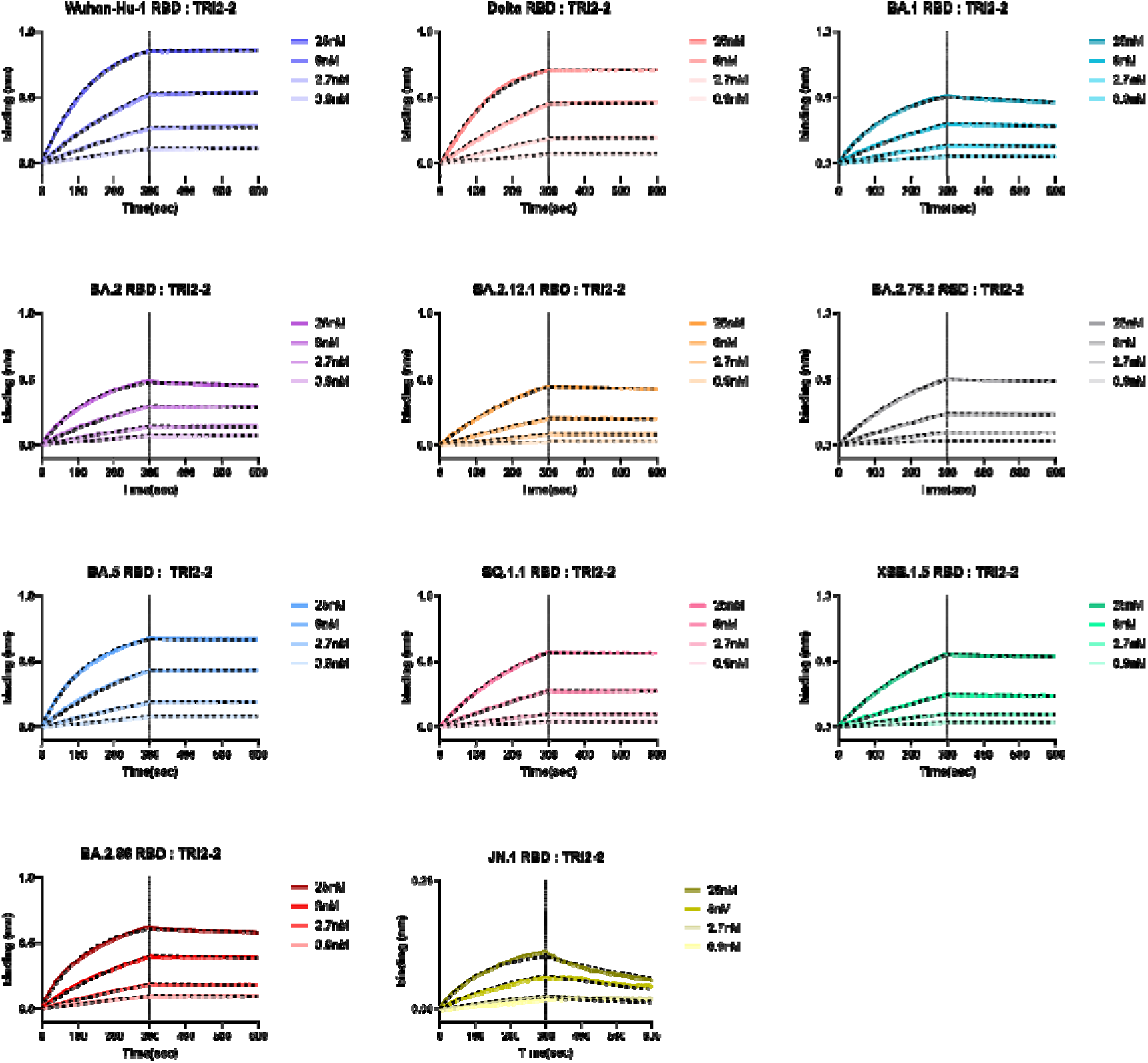
Kinetic analysis of TRI2-2 binding to SARS-CoV-2 variant RBDs immobilized on using biolayer interferometry. Biotinylated RBDs were immobilized on streptavidin biosensors to a final level of 1 nm shift each. The TRI2-2 concentrations used are provided in the color keys. Dashed black lines represent curve fits obtained using global fitting and a 1:1 binding model in the ForteBio BLI software. Representative graphs are shown from two biological replicates.

**Supplementary Figure 2.**
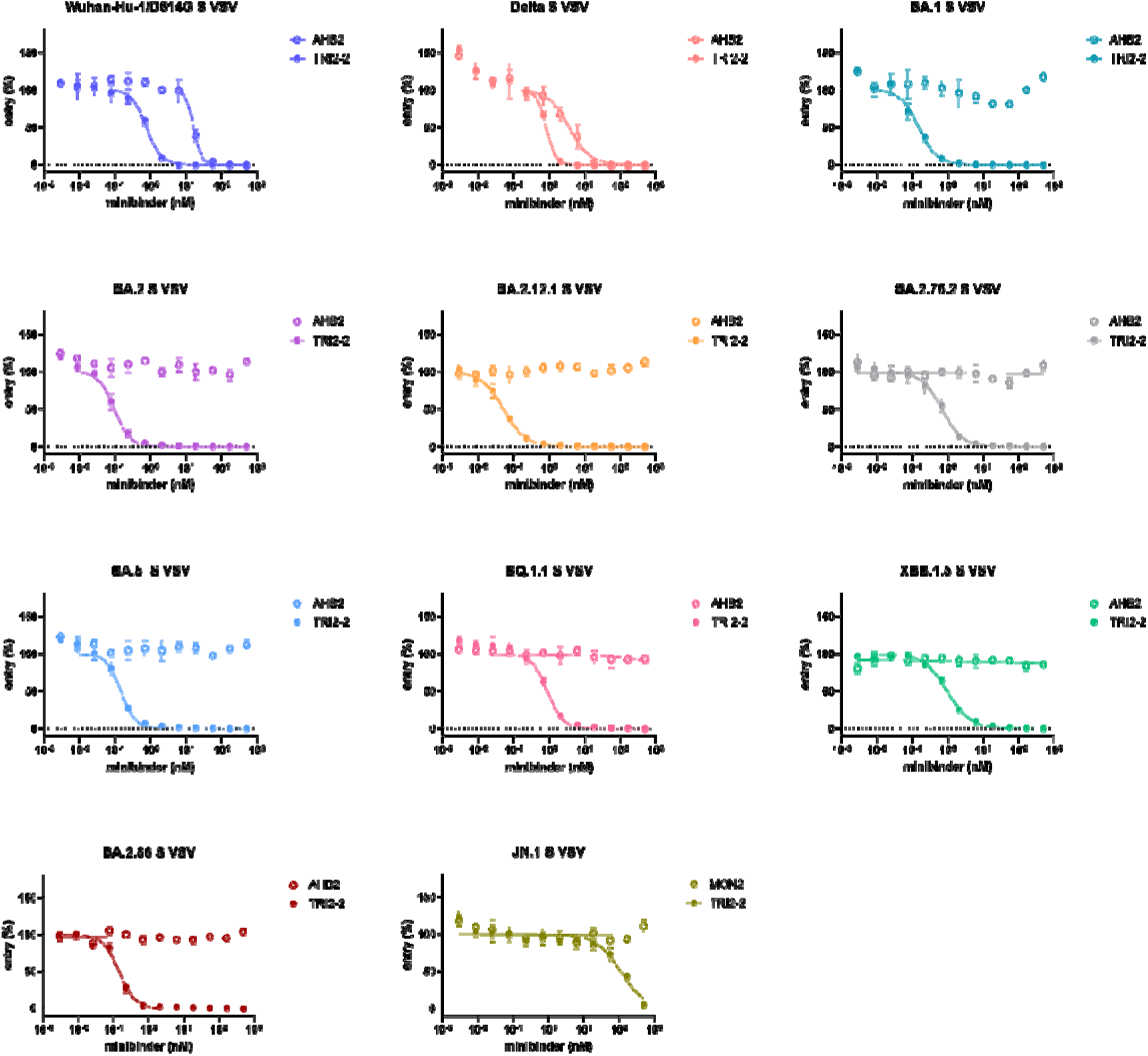
Dose-response curves for neutralization of SARS-CoV-2 S variant VSV pseudoviruses by the TRI2-2 and AHB2 miniprotein inhibitors. Each dot represents the mean of three technical replicates. SD shown as lines. Representative graphs are shown from three biological replicates.

**Supplementary Figure 3.**
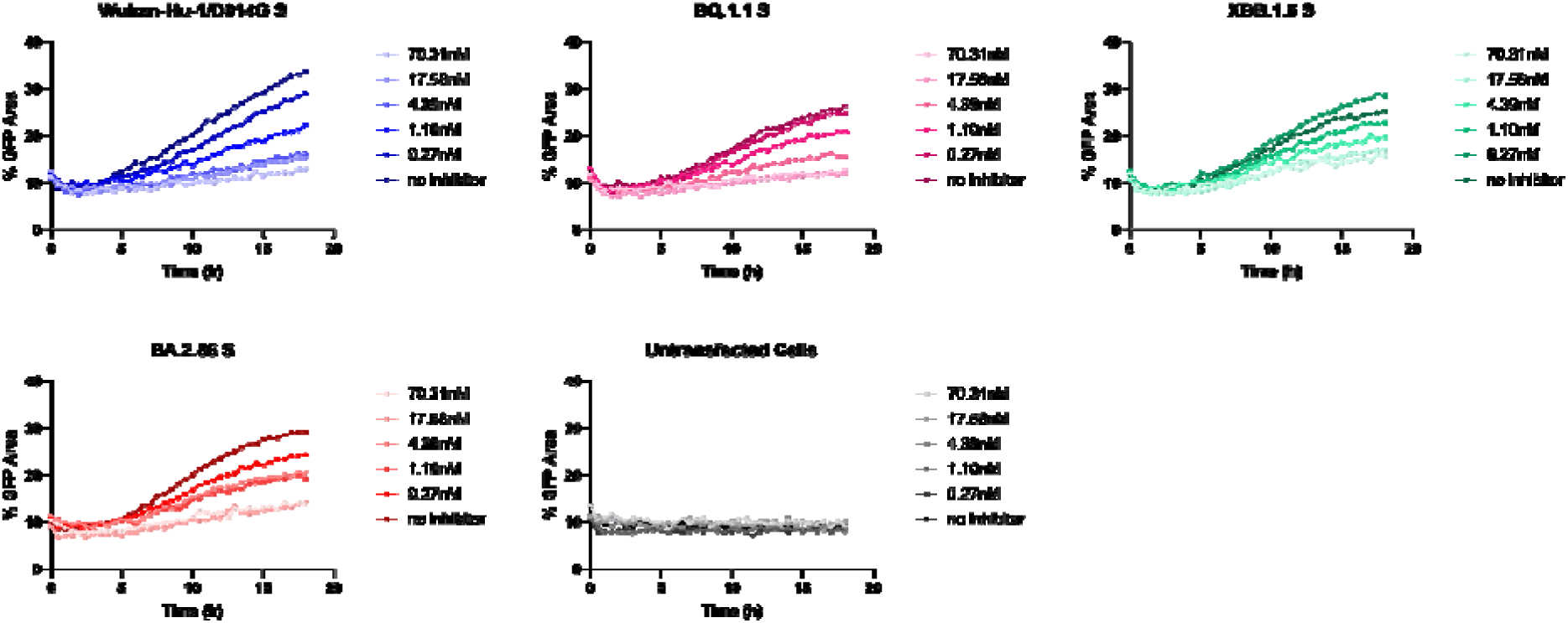
Dose-response curves for TRI2-2-mediated fusion inhibition of SARS-CoV-2 S variants. Representative graphs are shown from four biological replicates.

**Supplementary Figure 4.**
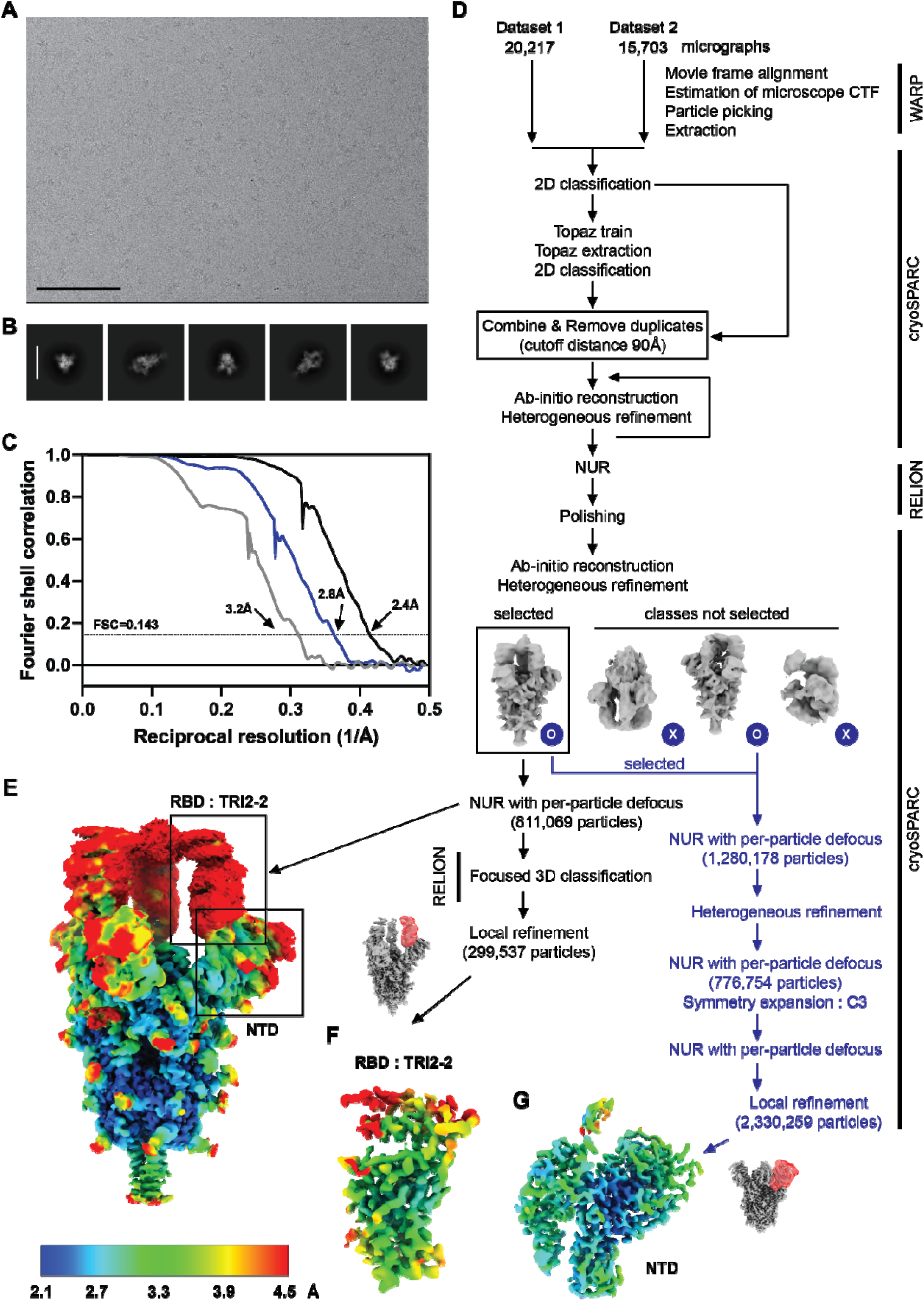
CryoEM data collection and refinement of TRI2-2 bound to the BA.2.86 S glycoprotein trimer. **(A-B)** Representative electron micrograph (A) and 2D class averages (B) of SARS-CoV-2 BA.2.86 S in complex with TRI2-2. The scale bar represents 100nm (A) and 210Å (B). **(C)** Gold-standard Fourier shell correlation curves for the cryoEM reconstructions. The 0.143 cutoff is indicated with a gray dashed line. Black, gray, and blue curves correspond to the global, RBD, and NTD reconstructions, respectively. **(D)** Data processing flowchart. NUR: non-uniform refinement. Masks used for local refinement are shown in red. **(E-G)** CryoEM map of SARS-CoV-2 BA.2.86 S in complex with TRI2-2 (E), locally refined map of the BA.2.86 RBD in complex with TRI2-(F), and locally refined map of the BA.2.86 NTD (G) colored by local resolution as determined using cryoSPARC.

**Supplementary Figure 5.**
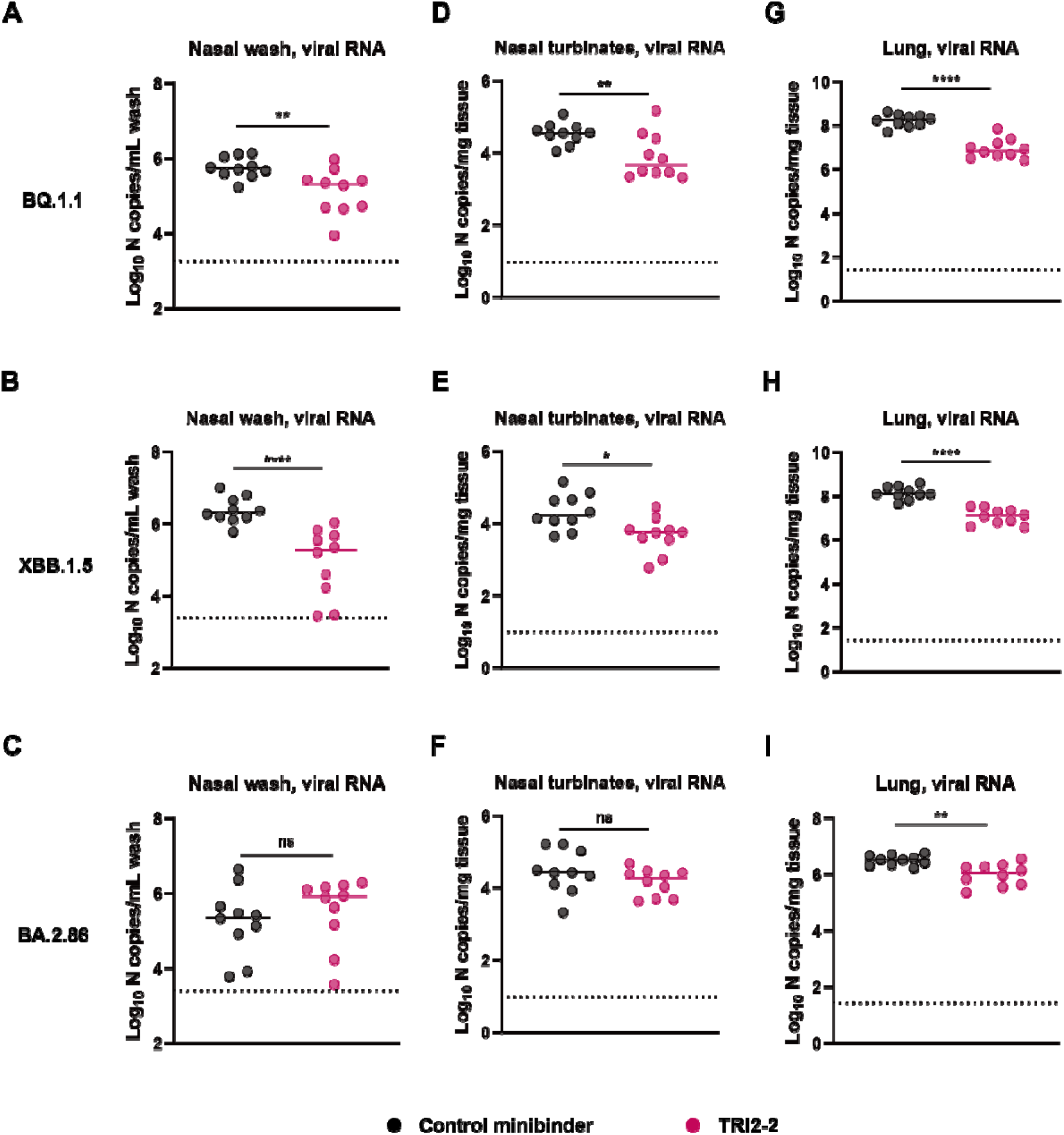
Quantification of viral RNA loads. **(A-C)** Genomic viral RNA levels in nasal washes for mice challenged with BQ.1.1 (A), XBB.1.5 (B), or BA.2.86 (C). **(D-F)** Genomic viral RNA levels in nasal turbinates for mice challenged with BQ.1.1 (D), XBB.1.5 (E), or BA.2.86 (F). **(G-I)** Genomic viral RNA levels in lungs for mice challenged with BQ.1.1 (G), or XBB.1.5 (H), or BA.2.86 (I) (lines indicate median; n□= 10 mice per group per virus challenge, two experiments; Two-tailed Mann-Whitney test between control and TRI2-2 treatment; ns, not significant; *P□<□0.05, **P<0.01, ***P□<□0.001, ****P□<□0.0001.).

**Supplementary Table 1.** Representative TRI2-2 binding kinetics and avidities (apparent affinities denoted K_D,app_) to SARS-CoV-2 variant RBDs obtained by biolayer interferometry. Values shown here are calculated from the curve fit from Supplementary Figure 1.

| | $K_{D,app}$ (M) | $K_{D,app}$ error | $k_{on}$ (1/Ms) | $k_{on}$ error | $k_{off}$ (1/s) | $k_{off}$ error |
| --- | --- | --- | --- | --- | --- | --- |
| **D614G** | 1.21E-12 | N/A | 2.86E+05 | 2.42E+02 | 3.46E-07 | N/A |
| **Delta** | 1.14E-12 | N/A | 2.59E+05 | 4.17E+02 | 2.95E-07 | N/A |
| **BA.1** | 9.28E-10 | 6.07E-12 | 2.84E+05 | 5.09E+02 | 2.64E-04 | 1.66E-06 |
| **BA.2** | 4.93E-10 | 5.82E-12 | 2.97E+05 | 5.47E+02 | 1.47E-04 | 1.71E-06 |
| **BA.2.12.1** | 8.51E-10 | 8.90E-12 | 1.37E+05 | 2.93E+02 | 1.17E-04 | 1.20E-06 |
| **BA.2.75.2** | 5.42E-10 | 6.40E-12 | 1.65E+05 | 2.67E+02 | 8.93E-05 | 1.05E-06 |
| **BA.5** | 3.06E-11 | 4.60E-12 | 3.25E+05 | 4.97E+02 | 9.92E-06 | 1.49E-06 |
| **BQ.1.1** | 2.10E-10 | 5.49E-12 | 1.60E+05 | 2.23E+02 | 3.36E-05 | 8.78E-07 |
| **XBB.1.5** | 6.58E-10 | 7.02E-12 | 1.60E+05 | 2.78E+02 | 1.05E-04 | 1.11E-06 |
| **BA.2.86** | 3.88E-10 | 5.79E-12 | 3.39E+05 | 6.69E+02 | 1.31E-04 | 1.95E-06 |
| **JN.1** | 6.19E-09 | 6.11E-11 | 2.82E+05 | 2.42E+03 | 1.75E-03 | 8.53E-06 |

**Supplementary Table 2.** CryoEM data collection and refinement statistics.

|  | SARS-CoV-2 S BA.2.86 in complex with minibinder TRI2-2 | SARS-CoV-2 BA.2.86 RBD in complex with TRI2-2 minibinder | SARS-CoV-2 BA.2.86 NTD |
| --- | --- | --- | --- |
| **Data collection and processing** | EMD-45972<br>PDB 9CWR | EMD-45969<br>PDB 9CWP | EMD-45971<br>PDB 9CWQ |
| Magnification | 105,000 | 105,000 | 105,000 |
| Voltage (kV) | 300 | 300 | 300 |
| Electron exposure (e<sup>-</sup>/Å<sup>2</sup>) | 53.25 | 53.25 | 53.25 |
| Defocus range (μm) | -0.8 - -1.8 | -0.8 - -1.8 | -0.8 - -1.8 |
| Pixel size (Å) | 0.829 | 0.829 | 0.829 |
| Symmetry imposed | C1 | C1 | C1 |
| Final particle images (no.) | 811,069 | 299,537 | 2,330,259 |
| Map resolution (Å) | 2.4 | 3.2 | 2.8 |
| FSC threshold | 0.143 | 0.143 | 0.143 |
| Map sharpening B factor (Å<sup>2</sup>) | -76.2 | -117.8 | -91.7 |
| Validation |  |  |  |
| MolProbity score | 1.02 | 0.82 | 1.64 |
| Clashscore | 1.80 | 0.79 | 2.33 |
| Poor rotamers (%) | 0.95 | 0.54 | 2.69 |
| Ramachandran plot |  |  |  |
| Favored (%) | 97.63 | 97.68 | 95.73 |
| Allowed (%) | 2.34 | 2.32 | 4.27 |
| Disallowed (%) | 0.03 | 0.00 | 0.00 |

## Methods

### Cells

Cell lines used in this study were DH10B competent cells (Thermo Fisher Scientific), HEK293T (ATCC, CRL-11268), Vero E6-TMPRSS2-GFP_11_ and BHK-21-GFP_1–10_ ^6^ and HEK293T cells with stable human ACE2 expression (kindly provided by Jesse Bloom)^13^. Cells were cultured in 10% FBS (Fisher Scientific-Cytiva), 1% penicillin-streptomycin (Thermo Fisher Scientific) DMEM at 37□, 5% CO_2_. BHK-21-GFP_1–10_ and Vero E6-TMPRSS2-GFP_11_ cells were generated in-house and were cultured supplemented with 2 µg/mL of Puromycin for BHK and 8 µg/mL of Puromycin and 4 µg/mL of Blasticidin for Vero cells in 10% FBS (Fisher Scientific-Cytiva), 1% penicillin-streptomycin (Thermo Fisher Scientific) DMEM at 37□, 5% CO_2_. Expi293F (Thermo Fisher Scientific) were cultured at 37°C and 8% CO_2_. None of the cell lines were authenticated or tested for mycoplasma contamination.

Vero-TMPRSS2^29^ cells were cultured at 37°C in Dulbecco’s modified Eagle medium (DMEM) supplemented with 10% fetal bovine serum (FBS), 10 □mM HEPES pH 7.3, 1 □mM sodium pyruvate, 1× non-essential-essential amino acids, and 100 □U/mL of penicillin–streptomycin. Vero-TMPRSS2 cells were supplemented with 5 µg/mL of blasticidin. Vero-hACE2-TMPRSS2 cells were supplemented with 10 µg/mL of puromycin. All cells routinely tested negative for mycoplasma using a PCR-based assay.

### Production of recombinant SARS-CoV-2 S RBDs

The SARS-CoV-2 RBDs were expressed in Expi293 cells (Thermo) at 37°C and 8% CO2. Cells were transfected with the corresponding plasmids using Expifectamine (Thermo) following the manufacturer’s protocol. Four days post-transfection, supernatants were clarified by centrifugation at 3000 g for 30 minutes, supplemented with 25 mM phosphate pH 8.0, and 300 mM NaCl and then passed through a 0.22µm sterile filter. Supernatants were then bound to 1 mL Histrap excel columns (Cytiva) previously equilibrated in 25 mM phosphate pH 8.0, 300 mM NaCl. Nickel columns were washed with 25 mM phosphate pH 8.0, 300 mM NaCl, and 10mM imidazole prior to elution with 25 mM phosphate pH 8.0, 300 mM NaCl and 300mM imidazole. After buffer exchanging into 20mM phosphate pH 8.0 and 100mM NaCl using a centrifugal filter device with a MWCO of 10kDa, the purified RBDs were biotinylated using the BirA biotin-protein ligase reaction kit (Avidity). The biotinylated RBD’s were bound, washed, and eluted again on the same affinity column. Purified biotinylated RBD’s were then concentrated and eluted on a Superdex200 increase 10/300 size-exclusion column (Cytiva) equilibrated in 20mM phosphate pH 8.0 and 100mM NaCl. Fractions containing monomeric and monodisperse RBDs were flash frozen and stored at −80°C until use.

### Production of recombinant TRI2-2 and influenza virus minibinders

The TRI2-2 and influenza virus minibinders were cloned into pET29b between the NdeI and XhoI restriction sites by Genscript. The TRI2-2 minibinder was cloned with a C-terminal polyhistidine tag and the influenza minibinder was cloned with an N-terminal polyhistidine tag^17^. Minibinders were expressed in Lemo21(DE3) cells (NEB) in TB II Media (MP Bio) at 37°C with IPTG induction. After cell harvest, pellets were resuspended in Gibco dPBS and lysed by microfluidization at 18,000 psi. Whole cell lysates were clarified by centrifugation at 18000 g for 30 minutes and supernatants were then bound to a 5 mL Histrap Nickel Sepharose FF column (Cytiva) previously equilibrated in Gibco dPBS supplemented with 30mM Imidazole. Nickel columns were washed with Gibco dPBS (ThermoFisher) supplemented with 30mM imidazole prior to elution with Gibco dPBS supplemented with 500mM Imidazole. Using a centrifugal filter device with a MWCO of 3kDa, the IMAC fractions containing minibinders of interest were concentrated and then further purified by size-exclusion chromatography using an Superdex S75 Increase 10/300 GL column (Cytiva) equilibrated in Gibco dPBS as running buffer. Fractions containing TRI2-2 or Influenza minibinders were further concentrated (as needed) filtered with a 0.2 μm filter, and then tested for endotoxin (LAL Charles River) prior to being flash frozen and stored at −80°C until use.

### Binding analysis using biolayer interferometry (BLI)

BLI binding assays were performed on an Octet Red (Sartorius) instrument operated at 30□ with shaking (1000 rpm). Streptavidin biosensors were hydrated in a 10x kinetics buffer (Sartorius) for 10 min prior to the experiment. Biosensors were incubated in a 10x kinetics buffer for 60s followed by the loading of biotinylated RBDs to the tip, all to a final level of 1 nm. Loaded biosensors were equilibrated in a 10x kinetics buffer for 120s. For affinity binding assays to determine K_D_ values, RBD-loaded tips were dipped into a concentration series of TRI2-2 (3-fold serial dilution from 25 nM to 0.9 nM) for 300 s followed by 300 s of dissociation in a 10x kinetics buffer. Global fits were used to calculate K_D_ values using a 1:1 binding fit model. Data were plotted using GraphPad Prism. Assays were replicated with two biological replicates (recombinant RBD proteins generated on different days) and representative graphs and values are shown in Supplementary Figure 1 and Supplementary Table 1, respectively.

### Production of VSV pseudoviruses

SARS-CoV-2 spike VSV pseudoviruses were produced using HEK293T cells seeded on BioCoat Cell Culture Dish: poly-D-Lysine 100 mm (Corning). The following day, cells were transfected with spike constructs using Lipofectamine 2000 (Thermo Fisher Scientific) in Opti-MEM transfection medium. After 5h of incubation at 37□°C with 5% CO2, cells were supplemented with DMEM containing 10% of FBS. On the next day, cells were washed with three times with DMEM and infected with VSV (G*ΔG-luciferase) for 2h, followed by five time wash with DMEM medium before addition of anti-VSV G antibody (I1-mouse hybridoma supernatant diluted 1:40, ATCC CRL-2700) and medium. After 18-24 h of incubation at 37□°C with 5% CO_2_, pseudoviruses were collected and cell debris removed by centrifugation at 3,000xg for 10 min. Pseudoviruses were further filtered using a 0.45 µm syringe filter and concentrated 10x prior to storage at −80°C.

### Neutralization assays

For SARS-CoV-2 S VSV neutralization with TRI2-2 and AHB2, HEK293T cells with stable human ACE2 expression in DMEM supplemented with 10% FBS and 1% PenStrep were seeded at 40,000 cells/well into 96-well plates [3610] (Corning) coated with poly-lysine [P4707] (Sigma) and incubated overnight at 37°C. The following day, a half-area 96-well plate (Greiner) was prepared with 3-fold serial dilutions of TRI2-2 and AHB2 with a starting concentration of 1 μM. An equal volume of DMEM with diluted pseudoviruses was added to each well. All pseudoviruses were diluted between 1:3 - 1:27 to reach a target entry of ∼10^6^ RLU. The mixture was incubated at room temperature for 45-60 minutes. Media was removed from the cells and 40 μL from each well of the half-area 96-well plate containing minibinder and pseudovirus were transferred to the 96-well plate seeded with cells and incubated at 37°C for 1h. After 1h, an additional 40 μL of DMEM supplemented with 20% FBS and 2% PenStrep was added to the cells. After 18–20h, 40 μL of One-Glo-EX substrate (Promega) was added to each well and incubated on a plate shaker in the dark for 5 min before reading the relative luciferase units using a BioTek Neo2 plate reader. Relative luciferase units were plotted, and normalized in Prism (GraphPad): 0% entry being cells lacking pseudovirus and 100% entry being cells containing virus but lacking minibinder. Prism (GraphPad) nonlinear regression with “[Inhibitor] versus normalized response with a variable slope” was used to determine IC_50_ values from curve fits with 3 technical repeats. 3 biological replicates were carried out for each sample-pseudovirus pair.

### Fusion assays

Cell to cell fusion assay using a split-GFP system was conducted as previously described. BHK-21-GFP_1–10_ cells were split into 6-well plates at a density of 250,000 cells per well. The following day, the growth medium was removed from the 6-well plates and cells were washed with DMEM followed by addition of the growth medium. Then, the cells were transfected with 4 µg of S protein DNA using Lipofectamine 2000 transfection kit. Vero E6-TMPRSS2-GFP_11_ cells were plated into 96-well, glass bottom, black-walled plates (CellVis) at a density of 36,000 cells per well. Twenty-four hours after transfection, BHK-21-GFP_1–10_ cells expressing the S protein were washed three times using FluoroBrite DMEM (Thermo Fisher) and detached using an enzyme-free cell dissociation buffer (Gibco). 9,000 BHK-21-GFP_1–10_ cells were added to each well with or without TRI2-2 with the 1:4 serial dilution starting from the initial concentration of 70nM, and the mixture was incubated at 37□°C and 5% CO_2_ for 2h before being transferred on top of the Vero E6-TMPRSS2-GFP_11_ that was washed 3 times with FluoroBrite DMEM. The mixture was incubated at 37□°C and 5% CO_2_ in a Cytation 7 plate Imager (BioTek) and both bright-field and GFP images were collected every 30□min for 18□h. Fusogenicity was assessed by measuring the area showing GFP fluorescence for each image using Gen5 Image Prime v3.11 software. Raw grayscale 16-bit images were pseudocolored in ImageJ using Green Hot Look Up Table.

### Production of recombinant SARS-CoV-2 S BA.2.86

The SARS-CoV-2 BA.2.86 hexapro S ectodomain construct includes its native signal peptide, hexapro mutations (F817P, A892P, A899, A942P, K986P, V987P), and a C-terminal foldon, avi tag, and a 8x histidine tag. SARS-CoV-2 BA.86 hexapro S ectodomain was expressed in Expi293 cells (Thermo) at 37°C and 8% CO_2_. Cells were transfected using Expifectamine293 (Thermo) following the manufacturer’s protocol. Four days post-transfection, Expi293 cell supernatant was clarified by centrifugation at 4,121g for 30 minutes, supplemented with 25 mM phosphate pH 8.0, 300 mM NaCl. Supernatant was then bound to His-Trap Excel column (Cytiva) previously equilibrated in 25 mM phosphate pH 8.0, 300 mM NaCl. Nickel columns were washed with 20-40mL of 25 mM phosphate pH 8.0, 300 mM NaCl, and 40mM Imidazole. S protein was eluted using 25 mM phosphate pH 8.0, 300 mM NaCl, and 300mM imidazole prior to being buffer exchanged to 50 mM Tris-HCl pH 8.0, 150 mM NaCl using a centrifugal filter device with a MWCO of 100 kDa. Protein was then flash frozen and stored at −80°C.

### Cryo-EM sample preparation and data collection

The SARS-CoV-2 BA.2.86 S complex with TRI2-2 at a molar ratio of 1:8 just before the grid preparation. The cryo-EM dataset was collected over two different sessions which were combined to be processed together. 3 µL of SARS-CoV-2 BA.2.86 S (Acro Biosystems, SPN-C524y) complex with TRI2-2 at 0.6mg/mL was added to a glow discharged (30s at 15mA) UltraAuFoil R1.2/1.3:Au300 grid^30^ prior to plunge freezing using a vitrobot MarkIV (ThermoFisher Scientific) with a blot force of −1, wait time of 10s, and 6 sec blot time at 100 % humidity and 22°C. 3.5 µL of SARS-CoV-2 BA.2.86 S produced following the aforementioned protein production complex with TRI2-2 at 0.2mg/mL was added to a glow discharged (10s at 15mA) Quantifoil 2nm C Au300 grid prior to plunge freezing using a vitrobot MarkIV (ThermoFisher Scientific) with a blot force of −1, 4 sec blot time, and 10s wait time at 100 % humidity and 22°C. Data were acquired using an FEI Titan Krios transmission electron microscope operated at 300 kV and equipped with a Gatan K3 direct detector and Gatan Quantum GIF energy filter, operated in zero-loss mode with a slit width of 20 eV. Automated data collection was carried out using serialEM^31^ at a nominal magnification of 105,000x with a pixel size of 0.829 Å. The dose rate was adjusted to 53e^−^/Å^2^, and each movie was acquired in counting mode fractionated in 79 frames of 50ms for UltraAuFoil and 99 frames of 40ms for Quantifoil dataset, respectively. A total of 20,217 and 15,703 micrographs were collected for each datasets, respectively. Stage was tilted 0, 30, and 45 degrees for collection with the UltraAuFoil grid.

### Cryo-EM data processing, model building and refinement

Particles were extracted with a box size of 320 pixels with a pixel size of 1.658 Å using WARP. Two rounds of reference-free 2D classification were performed using CryoSPARC^32^ to select well-defined particle images from each dataset. Particles belonging to classes with the best resolved spike protein density were selected. To improve particle picking further, we trained the Topaz^33^ picker on Warp-picked particle sets belonging to the selected classes after 2D classification. The particles picked using Topaz were extracted and subjected to 2D classification using cryoSPARC. The two different particle sets picked from Warp and Topaz were merged and duplicate particle picks were removed using a minimum distance cutoff of 90 Å. Initial model generation was done using ab-initio reconstruction in cryoSPARC and used as references for a heterogenous 3D refinement in cryoSPARC. After two rounds of ab-initio reconstructions and heterogeneous refinements to remove junk particles, 3D refinement was carried out using non-uniform refinement in cryoSPARC^34^ and the particles were transferred from cryoSPARC to Relion using pyem^35^ (https://github.com/asarnow/pyem) to be subjected to the Bayesian polishing procedure implemented in Relion^36^ during which particles were re-extracted with a box size of 512 pixels and a pixel size of 1.0 Å. After ab-initio reconstructions and heterogeneous refinements to select best class, subsequent 3D refinement used non-uniform refinement along with per-particle defocus refinement in cryoSPARC to yield the final reconstruction at 2.4 Aℒ resolution comprising 811,069 particles. To further improve the density at the RBD:TRI2-2 interface, 3D classification and local refinement was performed using Relion and cryoSPARC with a soft mask comprising the RBD and TRI2-2 yielding a reconstruction at 3.2□Å resolution enabling model building. Reported resolutions are based on the 0.143 gold-standard Fourier shell correlation (FSC) criterion and Fourier shell correlation curves were corrected for the effects of soft masking by high-resolution noise substitution^37,38^. To further improve the N terminus domain (NTD) density, particles belonging to the best selected classes were subjected to another round of heterogeneous refinement, followed by non-uniform refinement with per-particle defocus. Particles were then symmetry expanded following C3 axis and local refinement was performed using cryoSPARC with a soft mask comprising the NTD domain yielding a reconstruction at 2.8□Å resolution enabling model building. UCSF Chimera, UCSF ChimeraX, and Coot were used to fit and rebuild atomic models into the cryoEM maps utilizing sharpened and unsharpened maps. The models were refined and relaxed using Rosetta^39,40^ and validated using Phenix^41^, Molprobity^42^ and Privateer^43^.

### Virus

The BQ.1.1 (hCoV-19/USA/CA-Stanford-106_S04/2022; EPI_ISL_15196219) and XBB.1.5 hCoV-19/USA/MD-HP40900-PIDYSWHNUB/2022; EPI_ISL_16026423 strains were obtained from nasopharyngeal isolates and provided as generous gifts by Mehul Suthar (Emory University) and Andrew Pekosz (Johns Hopkins), respectively. All virus stocks were generated in Vero-TMPRSS2 cells and subjected to next-generation sequencing as described previously^29^ to confirm the presence and stability of expected substitutions. All virus experiments were performed in an approved biosafety level 3 (BSL-3) facility.

Mouse experiments Animal studies were carried out in accordance with the recommendations in the Guide for the Care and Use of Laboratory Animals of the National Institutes of Health. The protocols were approved by the Institutional Animal Care and Use Committee at the Washington University School of Medicine (assurance number A3381-01). Virus inoculations were performed under anesthesia that was induced and maintained with ketamine hydrochloride and xylazine, and all efforts were made to minimize animal suffering.

Heterozygous K18-hACE2 C57BL/6J mice (strain: 2B6.Cg-Tg(K18-ACE2)2Prlmn/J) were obtained from The Jackson Laboratory. All animals were housed in groups and fed standard chow diets. For mouse experiments, 8-week-old female K18-hACE2 mice were administered 10^4^ FFU of the respective SARS-CoV-2 strains by intranasal administration. One day later, animals were administered a single 10 mg/kg dose of influenza-specific control or TRI2-2 minibinder intranasally. *In vivo* studies were not blinded, and mice were randomly assigned to treatment groups. No sample-size calculations were performed to power each study. Instead, sample sizes were determined based on prior *in vivo* virus challenge experiments.

### Measurement of viral RNA levels

Tissues were weighed and homogenized with zirconia beads in a MagNA Lyser instrument (Roche Life Science) in 1 mL of DMEM medium supplemented with 2% heat-inactivated FBS. Tissue homogenates were clarified by centrifugation at 10,000 rpm for 5 min and stored at −80°C. RNA was extracted using the MagMax mirVana Total RNA isolation kit (Thermo Fisher Scientific) on the Kingfisher Flex extraction robot (Thermo Fisher Scientific). RNA was reverse transcribed and amplified using the TaqMan RNA-to-CT 1-Step Kit (Thermo Fisher Scientific). Reverse transcription was carried out at 48°C for 15 min followed by 2 min at 95°C. Amplification was accomplished over 50 cycles as follows: 95°C for 15 s and 60°C for 1 min. Copies of SARS-CoV-2 N gene RNA in samples were determined using a previously published assay^44^. Briefly, a TaqMan assay was designed to target a highly conserved region of the N gene (Forward primer: ATGCTGCAATCGTGCTACAA; Reverse primer: GACTGCCGCCTCTGCTC; Probe: /56-FAM/TCAAGGAAC/ZEN/AACATTGCCAA/3IABkFQ/). This region was included in an RNA standard to allow for copy number determination down to 10 copies per reaction. The reaction mixture contained final concentrations of primers and probes of 500 and 100 nM, respectively.

### Viral plaque assay

Vero-TMPRSS2-hACE2 cells were seeded at a density of 1 × 10^5^ cells per well in 24-well tissue culture plates. The following day, medium was removed and replaced with 200 µL of material to be titrated diluted serially in DMEM supplemented with 2% FBS. After 1 h, 1 mL of methylcellulose overlay was added. Plates were incubated for 72 h, then fixed with 4% paraformaldehyde (final concentration) in phosphate-buffered saline (PBS) for 20 min. Plates were stained with 0.05% (wt/vol) crystal violet in 20% methanol and washed twice with distilled, deionized water.

### Statistical analysis

All statistical tests were performed as described in the indicated figure legends using Prism 9.4.1 or 10.1.1. When comparing against control value in fusion assay, one sample t test was performed to determine statistical significance. When comparing two groups in viral challenge studies, a Mann-Whitney test was performed to determine statistical significance. The number of independent experiments performed is indicated in the relevant figure legends.

### Data availability

The sharpened and unsharpened cryoEM reconstructions and atomic models of SARS-CoV-2 BA.2.86 S in complex with TRI2-2 minibinder, SARS-CoV-2 BA.2.86 RBD in complex with TRI2-2 minibinder, and SARS-CoV-2 BA.2.86 NTD have been deposited in the Electron Microscopy Data Bank and the Protein Data Bank with accession codes EMD-45972 and PDB 9CWR (SARS-CoV-2 BA.2.86 S in complex with TRI2-2 minibinder), EMD-45969 and PDB 9CWP (SARS-CoV-2 BA.2.86 RBD in complex with TRI2-2 minibinder), and EMD-45971 and PDB 9CWQ (SARS-CoV-2 BA.2.86 NTD). Other data will be available from the corresponding author upon request.

## References

1. Walls, A. C. et al. Structure, Function, and Antigenicity of the SARS-CoV-2 Spike Glycoprotein. Cell 181, 281–292.e6 (2020).

2. Zhou, P. et al. A pneumonia outbreak associated with a new coronavirus of probable bat origin. Nature (2020) doi:10.1038/s41586-020-2012-7.

3. Hoffmann, M. et al. SARS-CoV-2 Cell Entry Depends on ACE2 and TMPRSS2 and Is Blocked by a Clinically Proven Protease Inhibitor. Cell 181, 271–280.e8 (2020).

4. Letko, M., Marzi, A. & Munster, V. Functional assessment of cell entry and receptor usage for SARS-CoV-2 and other lineage B betacoronaviruses. Nat Microbiol 5, 562–569 (2020).

5. Walls, A. C. et al. SARS-CoV-2 breakthrough infections elicit potent, broad, and durable neutralizing antibody responses. Cell (2022) doi:10.1016/j.cell.2022.01.011.

6. Bowen, J. E. et al. Omicron spike function and neutralizing activity elicited by a comprehensive panel of vaccines. Science eabq0203 (2022).

7. Cameroni, E. et al. Broadly neutralizing antibodies overcome SARS-CoV-2 Omicron antigenic shift. Nature 602, 664–670 (2022).

8. Viana, R. et al. Rapid epidemic expansion of the SARS-CoV-2 Omicron variant in southern Africa. Nature (2022) doi:10.1038/d41586-021-03832-5.

9. Tegally, H. et al. Emergence of SARS-CoV-2 Omicron lineages BA.4 and BA.5 in South Africa. Nat. Med. 28, 1785–1790 (2022).

10. Cao, Y. et al. Imprinted SARS-CoV-2 humoral immunity induces convergent Omicron RBD evolution. Nature (2022) doi:10.1038/s41586-022-05644-7.

11. Cao, L. et al. De novo design of picomolar SARS-CoV-2 miniprotein inhibitors. Science 370, 426–431 (2020).

12. Hunt, A. C. et al. Multivalent designed proteins neutralize SARS-CoV-2 variants of concern and confer protection against infection in mice. Sci. Transl. Med. 0, eabn1252.

13. Crawford, K. H. D. et al. Protocol and Reagents for Pseudotyping Lentiviral Particles with SARS-CoV-2 Spike Protein for Neutralization Assays. Viruses 12, (2020).

14. Kodaka, M. et al. A new cell-based assay to evaluate myogenesis in mouse myoblast C2C12 cells. Exp. Cell Res. 336, 171–181 (2015).

15. Bussani, R. et al. Persistence of viral RNA, pneumocyte syncytia and thrombosis are hallmarks of advanced COVID-19 pathology. EBioMedicine 61, 103104 (2020).

16. Winkler, E. S. et al. SARS-CoV-2 infection of human ACE2-transgenic mice causes severe lung inflammation and impaired function. Nat. Immunol. 21, 1327–1335 (2020).

17. Cao, L. et al. Design of protein-binding proteins from the target structure alone. Nature 605, 551–560 (2022).

18. Robert, X. & Gouet, P. Deciphering key features in protein structures with the new ENDscript server. Nucleic Acids Res. 42, W320–4 (2014).

19. Park, Y.-J. et al. Imprinted antibody responses against SARS-CoV-2 Omicron sublineages. Science eadc9127 (2022).

20. Tang, J., et al. Respiratory mucosal immunity against SARS-CoV-2 after mRNA vaccination. Sci Immunol 7, eadd4853 (2022).

21. Yisimayi, A. et al. Repeated Omicron exposures override ancestral SARS-CoV-2 immune imprinting. Nature 625, 148–156 (2024).

22. Mao, T. et al. Unadjuvanted intranasal spike vaccine elicits protective mucosal immunity against sarbecoviruses. Science 378, eabo2523 (2022).

23. Oh, J. E. et al. Intranasal priming induces local lung-resident B cell populations that secrete protective mucosal antiviral IgA. Science Immunology 6, eabj5129 (2021).

24. Langel, S. N. et al. Adenovirus type 5 SARS-CoV-2 vaccines delivered orally or intranasally reduced disease severity and transmission in a hamster model. Sci. Transl. Med. eabn6868 (2022).

25. Hassan, A. O. et al. A single intranasal dose of chimpanzee adenovirus-vectored vaccine protects against SARS-CoV-2 infection in rhesus macaques. Cell Rep Med 2, 100230 (2021).

26. Ying, B. et al. Author Correction: Mucosal vaccine-induced cross-reactive CD8+ T cells protect against SARS-CoV-2 XBB.1.5 respiratory tract infection. Nat. Immunol. 25, 578 (2024).

27. de Vries, R. D. et al. Intranasal fusion inhibitory lipopeptide prevents direct-contact SARS-CoV-2 transmission in ferrets. Science 371, 1379–1382 (2021).

28. Song, R. et al. Post-exposure prophylaxis with SA58 (anti-SARS-COV-2 monoclonal antibody) nasal spray for the prevention of symptomatic COVID-19 in healthy adult workers: a randomized, single-blind, placebo-controlled clinical study. Emerg. Microbes Infect. 12, 2212806 (2023).

29. Chen, R. E. et al. Resistance of SARS-CoV-2 variants to neutralization by monoclonal and serum-derived polyclonal antibodies. Nat. Med. 27, 717–726 (2021).

30. Russo, C. J. & Passmore, L. A. Electron microscopy: Ultrastable gold substrates for electron cryomicroscopy. Science 346, 1377–1380 (2014).

31. Mastronarde, D. N. Automated electron microscope tomography using robust prediction of specimen movements. J. Struct. Biol. 152, 36–51 (2005).

32. Punjani, A., Rubinstein, J. L., Fleet, D. J. & Brubaker, M. A. cryoSPARC: algorithms for rapid unsupervised cryo-EM structure determination. Nat. Methods 14, 290–296 (2017).

33. Bepler, T. et al. Positive-unlabeled convolutional neural networks for particle picking in cryo-electron micrographs. Nat. Methods 16, 1153–1160 (2019).

34. Punjani, A., Zhang, H. & Fleet, D. J. Non-uniform refinement: adaptive regularization improves single-particle cryo-EM reconstruction. Nat. Methods 17, 1214–1221 (2020).

35. Asarnow, D., Palovcak, E. & Cheng, Y. UCSF pyem v0. 5. Zenodo 10.5281/zenodo 3576630, 2019 (2019).

36. Zivanov, J., Nakane, T. & Scheres, S. H. W. A Bayesian approach to beam-induced motion correction in cryo-EM single-particle analysis. IUCrJ 6, 5–17 (2019).

37. Rosenthal, P. B. & Henderson, R. Optimal determination of particle orientation, absolute hand, and contrast loss in single-particle electron cryomicroscopy. J. Mol. Biol. 333, 721– 745 (2003).

38. Chen, S. et al. High-resolution noise substitution to measure overfitting and validate resolution in 3D structure determination by single particle electron cryomicroscopy. Ultramicroscopy 135, 24–35 (2013).

39. Frenz, B. et al. Automatically Fixing Errors in Glycoprotein Structures with Rosetta. Structure 27, 134–139.e3 (2019).

40. Wang, R. Y. et al. Automated structure refinement of macromolecular assemblies from cryo-EM maps using Rosetta. Elife 5, (2016).

41. Liebschner, D. et al. Macromolecular structure determination using X-rays, neutrons and electrons: recent developments in Phenix. Acta Crystallogr D Struct Biol 75, 861–877 (2019).

42. Chen, V. B. et al. MolProbity: all-atom structure validation for macromolecular crystallography. Acta Crystallogr. D Biol. Crystallogr. 66, 12–21 (2010).

43. Agirre, J. et al. Privateer: software for the conformational validation of carbohydrate structures. Nat. Struct. Mol. Biol. 22, 833–834 (2015).

44. Case, J. B., Bailey, A. L., Kim, A. S., Chen, R. E. & Diamond, M. S. Growth, detection, quantification, and inactivation of SARS-CoV-2. Virology 548, 39–48 (2020).

